# Mitotic systemic genomic instability in yeast

**DOI:** 10.1101/161869

**Authors:** Nadia M. V. Sampaio, Aline Rodrigues-Prause, V. P. Ajith, Theodore M. Gurol, Mary J. Chapman, Ewa P. Malc, Parijat Chakraborty, Fabiana M. Duarte, Guadalupe M. Aguirre, Pedro A. Tizei, Gonçalo A. G. Pereira, Piotr A. Mieczkowski, Koodali T. Nishant, Juan Lucas Argueso

## Abstract

Conventional models of genome evolution generally include the assumption that mutations accumulate gradually and independently over time. We characterized the occurrence of sudden spikes in the accumulation of genome-wide loss-of-heterozygosity (LOH) in *Saccharomyces cerevisiae*, suggesting the existence of a mitotic systemic genomic instability process (mitSGI). We characterized the emergence of a rough colony morphology phenotype resulting from an LOH event spanning a specific locus (*ACE2/ace2-A7*). Surprisingly, half of the clones analyzed also carried unselected secondary LOH tracts elsewhere in their genomes. The number of secondary LOH tracts detected was 20-fold higher than expected assuming independence between mutational events. Secondary LOH tracts were not detected in control clones without a primary selected LOH event. We then measured the rates of single and double LOH at different chromosome pairs and found that coincident LOH accumulated at rates 30-100 fold higher than expected if the two underlying single LOH events occurred independently. These results were consistent between two different strain backgrounds, and in mutant strains incapable of entering meiosis. Our results indicate that a subset of mitotic cells within a population experience systemic genomic instability episodes, resulting in multiple chromosomal rearrangements over one or few generations. They are reminiscent of early reports from the classic yeast genetics literature, as well as recent studies in humans, both in the cancer and genomic disorder contexts, all of which challenge the idea of gradual accumulation of structural genomic variation. Our experimental approach provides a model to further dissect the fundamental mechanisms responsible for mitSGI.

**SIGNIFICANCE STATEMENT:** Point mutations and alterations in chromosome structure are generally thought to accumulate gradually and independently over many generations. Here, we combined complementary genetic approaches in budding yeast to track the appearance of chromosomal changes resulting in loss-of-heterozygosity (LOH). Contrary to expectations, our results provided evidence for the occurrence of non-independent accumulation of multiple LOH events over one or a few cell generations. These results are analogous to recent reports of bursts of chromosomal instability in humans. Our experimental approach provides a framework to further dissect the fundamental mechanisms underlying systemic chromosomal instability processes, including in the human cancer and genomic disorder contexts.

## INTRODUCTION

Heterozygosity is often associated with beneficial phenotypes in a variety of multicellular eukaryotes ranging from plants, to livestock, and even humans (1). At the organismal level, heterozygosity can be promoted and maintained through breeding between unrelated individuals, and conversely, can be lost through inbreeding (2). It can also be lost at the cellular level through allelic mitotic recombination between homologous chromosomes. Such loss-of-heterozygosity (LOH) events typically have negative consequences, such as somatic mosaicism or loss of tumor suppressor genes (3), but unless these mutations occur in the germline, they are not heritable and do not have long term consequences for the species.

Single cell eukaryotes including various yeast species also benefit from heterozygous genomes (4, 5). However, maintaining heterozygosity is more challenging in these cases as a mitotic LOH event leads to immediate fixation of the homozygous state in a clonal cell lineage. High levels of genomic heterozygosity have been described in several *Saccharomyces cerevisiae* strains (6-8). One of the first examples to be characterized was the JAY270/PE-2 strain used in bioethanol production (6). This heterothallic diploid was originally isolated as a robust and highly productive contaminant at a sugarcane distillery (9). Similarly isolated wild strains are also heterothallic and heterozygous (10), and genomic heterozygosity is suspected to contribute to their industrial traits. Interestingly, in most of the strains described above, including JAY270/PE-2, heterozygosity is not evenly distributed across the genome. Heterozygous regions are interspersed with stretches of homozygosity indicating the occurrence of LOH events in clonal ancestors (7). However, it is still unclear what, if any, consequences these LOH events may have on their general fitness.

The relationship between genomic heterozygosity, LOH, and phenotypic consequences in yeast is better understood in the human pathogen *Candida albicans*. In that system, LOH events have been shown to have a profound effect on clinically relevant traits, particularly drug resistance (5, 11). For example, LOH leading to homozygosis of a hyperactive form of Tac1, a transcription factor that regulates the multidrug transporter genes, results in increased efflux of azole antifungal drugs (12). In addition to providing a recurrent path to drug resistance, LOH has been shown to play a significant role in the evolution of the *C. albicans* genome (13, 14).

In this study, we identified and characterized a specific and easily discernible phenotypic transition in the *S. cerevisiae* JAY270/PE-2 strain, from smooth to rough yeast colony morphology, caused by an LOH event spanning the *ACE2* locus on chromosome XII (Chr12). Whole genome analyses of rough clones selected for carrying this specific Chr12 LOH event revealed that additional unselected recombination events were often present elsewhere in the genome. This initial observation was validated by direct measurements of coincident LOH rates at different chromosomes, suggesting the existence of a mitotic systemic genomic instability (mitSGI) process. The high rate of coincident LOH uncovered in our study resembles the bursts of accumulation of copy number alterations (CNAs) in human cancer (15) and genomic disorders (16). The results reported here have important ramifications for the characterization of mitSGI mechanisms that contribute to structural genomic variation.

## RESULTS

### Appearance of altered colony morphology derivatives of JAY270

One of the most desirable features of the JAY270/PE-2 bioethanol production strain (henceforth referred to simply as JAY270) is that it does not normally aggregate during industrial sugarcane extract fermentation (*i.e.* cells stay in suspension in liquid culture). Accordingly, JAY270 produces normal hemispherical colonies with smooth surfaces and edges when grown in solid agar medium (Fig. 1A). While this is the phenotype typically observed, over the course of our studies using this strain we noticed the sporadic occurrence of colonies clonally derived from JAY270 that displayed altered morphology: relatively flat-growing colonies with rough surfaces and edges (Fig. 1B). Under bright field microscopic examination, yeast cells derived from such rough colonies appeared to grow in chains, showing a budding pattern consistent with a defect in the separation of the daughter cells from their mother (Fig. 1C-D). We stained these cells with calcofluor white to visualize the chitin-rich ring septa, confirming the attachment of mother and daughter cells at the budding neck site (Fig. 1E-H).

**Figure 1.**
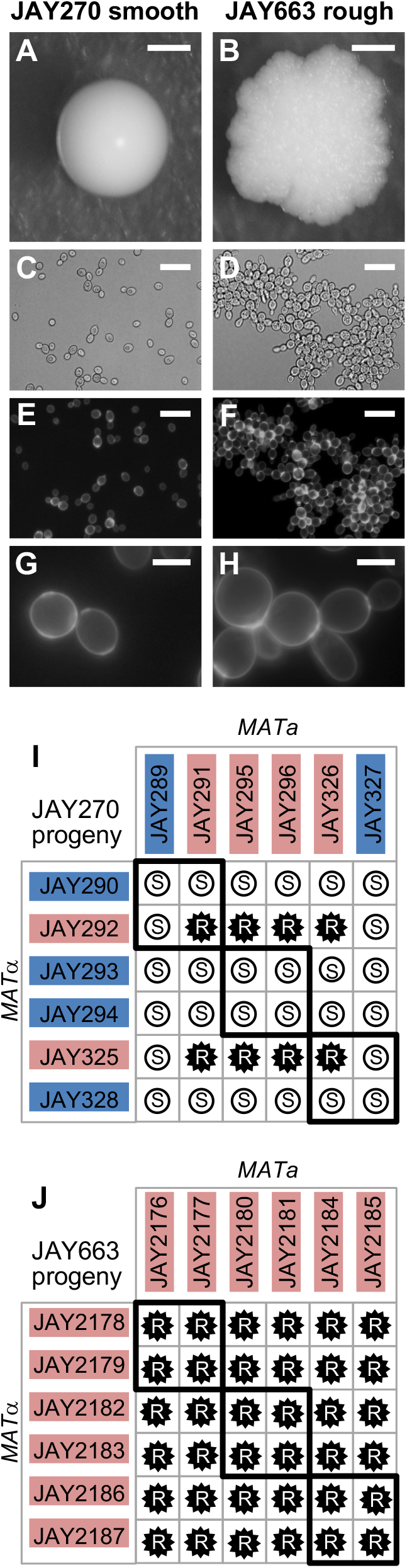
Smooth and rough colony morphologies, mother-daughter cell attachment, and phenotypes of diploids derived from mating specific haploids. **A-H** show images of the JAY270 smooth parent diploid strain (left panels) and its spontaneous rough derivative JAY663 (right panels). **A** and **B**, colony morphologies on YPD agar after 3 days growth at 30C. **C** and **D**, bright field, and **E-H**, fluorescence microscopy of cells stained with calcofluor white to highlight chitin septa and the mother-daughter cell attachment. Scale bars are 1mm (**A-B**), 20µm (**C-F**) and 5µm (**G-H**). **I** and **J**, Smooth (**S**, white circles) and rough (**R**, black stars) phenotypes of diploids formed by crossing the indicated *MATa* and *MATα* haploids isolated from three tetrads of each JAY270 (**I**) and JAY663 (**J**). Thick black lines indicate the four diploids derived from matings of intra-tetrad sibling haploids. The colored backgrounds for each haploid correspond to their inferred genotype (Blue, dominant wild type allele; Red, recessive mutant allele). All 12 haploids from panel **I** had their whole genomes sequenced. Co segregation analysis with JAY270 HetSNPs (Fig. S1) was used for identification of the causal mutation at the *ACE2* locus (Figs. S2 and S3).

We initially isolated five independent examples of such rough colonies for genetic characterization (JAY663, JAY664, JAY665, JAY912 and JAY913), all of which were derived either directly from JAY270 or from JAY270-isogenic strains. The phenotype of these isolates was stably maintained and was not reversible over several clonal generations, suggesting that it was likely hard-wired genetically and not caused by a transient transcriptional or post-transcriptional state. We estimated that these five rough colony isolates appeared spontaneously from a pool of ~50,000 smooth colonies. Assuming a genetic origin and based on this high frequency of occurrence in diploid cells, we reasoned that this phenotype was unlikely to be caused by a rare dominant *de novo* nucleotide point mutation, but instead, mitotic recombination leading to loss-of-heterozygosity (LOH) provided a more plausible mechanism.

In a parallel project, we observed that crossing two specific haploid descendants of JAY270 (JAY291 *MATa* and JAY292 *MATa*) resulted in diploid cells with the same rough colony morphology and mother-daughter cell attachment pattern observed in the five rough-colony isolates above. This was despite the fact that JAY291, JAY292, and all other haploid derivatives of JAY270 have the normal smooth colony phenotype. This indicated that the rough colony phenotype was diploid-specific, and the ability to consistently reproduce the mutant phenotype in controlled crosses between specific haploids opened an avenue to investigate its genetic basis.

We previously reported the whole genome sequence of the JAY291 haploid (6). Since then we have sequenced the genomes of 55 additional JAY270-derived haploids. This genome sequence dataset, comprising fourteen sets of four-spore tetrads, was generated in a project to characterize the abundance, distribution and phasing of heterozygous loci in the JAY270 genome, the full results of which will be described elsewhere. These haploid genomic sequences were used to create a draft map of phased heterozygous single nucleotide polymorphisms (HetSNPs) containing 12,023 loci unevenly distributed across the genome (Fig. S1). We carried out crosses between twelve sequenced haploid descendants of JAY270 (3 tetrads; Fig. 1I). All possible *MATa* x *MATa* crosses were performed producing 36 different diploids. Among them, we found eight with rough and 28 with smooth colony surfaces, in a pattern that was consistent with recessive inheritance of a trait controlled by a single gene. Even though the rough colony phenotype was not observed in any of the haploid parents, the phenotypes of their respective diploid combinations allowed us to infer which allele was present in the parents: either the wild type dominant allele or the recessive mutant allele.

In addition, we induced sporulation of one of the spontaneous rough-colony isolates, JAY663, dissected tetrads, and examined the phenotypes of the haploid derivatives. None of the resulting haploids displayed the rough colony phenotype; they were all smooth (~100 examined). We then took twelve of these haploids (JAY2176 through JAY2187 comprising three full tetrads, determined their mating types, and conducted all possible mating combinations between them (Fig. 1J). In this case, all 36 crosses resulted in rough colony diploids. This result was consistent with JAY663 being homozygous for the causal recessive mutant allele, and supported the hypothesis that copy neutral LOH could be responsible for the sporadic appearance of the mutant phenotype in JAY270.

### Genetic basis of the rough colony phenotype

Based on the interpretation that the rough colony phenotype was associated with monogenic recessive inheritance of a diploid-specific trait, we divided the sequenced JAY270-derived haploids from Fig. 1I into two groups according to their inferred genotype. Group 1 included the six haploids inferred to carry the mutant recessive allele, whereas group 2 included the six haploids with the wild type dominant allele. We then compared the genome sequences of the twelve haploids to the draft JAY270 HetSNPs map. We interrogated each of the HetSNPs searching for alleles that cosegregated in all six individuals within group 1, and that conversely, had the other allele co-segregating in all six individuals within group 2. This analysis identified two candidate regions that fit the strict co-segregation criterion (Fig. S2A-C). One of the candidate regions corresponded to ~30 Kb on Chr11, including thirteen genes; and the other spanned ~15 Kb containing nine genes on the right arm of Chr12, located ~50 Kb centromere proximal to the ribosomal DNA genes tandem repeats (rDNA).

We reviewed the annotations of the 22 candidate genes, and identified a gene located in the Chr12 region, *ACE2*, which encodes a transcription factor that controls the expression of genes involved in the mother-daughter cell separation process (17). In cells lacking Ace2p, the daughter cell remains attached to the mother cell wall at the bud neck, resulting in the accumulation of multicellular clusters. Importantly, a diploid-specific rough colony phenotype is observed in *ace2*/*ace2* homozygous mutant strains in certain genetic backgrounds (18).

We inspected the genomic sequence of the *ACE2* gene in JAY291, and compared it to the sequence in the S288c reference genome. Only one difference was identified: The wild type *ACE2* allele in S288c contains a homopolymer run of eight adenine nucleotides, while the mutant allele in JAY291 has seven adenines in this region, resulting in a -1 frameshift mutation and a stop codon shortly downstream. Hence, we named the mutant allele *ace2-A7*. We then conducted reciprocal complementation tests to formally demonstrate that *ace2-A7* was the causal mutation responsible for the rough colony phenotype. The mutant alleles in haploids JAY291 and JAY292 were replaced with the wild type allele, resulting respectively in the isogenic *ACE2* strains JAY1051 and JAY1039. When these allele replacement strains were crossed to *ace2-A7* strains (Figure S2D), the resulting diploids displayed the smooth colony phenotype, thus confirming that the wild type *ACE2* allele fully complemented the *ace2-A7* mutation in heterozygous diploids.

### Analysis of Chr12 LOH in spontaneous rough colony isolates

After identifying the association between the *ACE2* locus and the rough colony phenotype, we determined its sequence in the five spontaneous rough colony derivatives isolated earlier in the study. We PCR-amplified and Sanger-sequenced the region containing the adenine homopolymer run in *ACE2* from JAY270, from the haploid derivatives JAY290 and JAY291, and from the rough colony isolates (Fig S3A-B). This analysis confirmed the presence of a run of 8 adenines in *ACE2* (JAY290) and 7 adenines in *ace2-A7* (JAY291). The chromatogram in the JAY270 heterozygous diploid was consistent with a mixture of *ACE2* and *ace2-A7* DNA templates being present in the sequencing reaction: single nucleotide peaks were observed at positions primer-proximal to the homopolymer run, and out-of-register double peaks were seen downstream of the seventh adenine nucleotide. The chromatograms for all five rough colony isolates showed the presence of the *ace2-A7* frameshift mutation and absence of the *ACE2* allele. The loss of the *ACE2* allele in the diploid rough-colony isolates may be explained by either a copy-neutral LOH mechanism such as inter-homolog mitotic recombination, by a segmental deletion spanning *ACE2*, or Chr12 monosomy. To distinguish between these scenarios, we conducted tetrad analysis with the five spontaneous rough colony isolates. Four of them produced tetrads that had four viable haploid spores, and each spore had a copy of the *ACE2* locus as determined by PCR (data not shown). One of the isolates, JAY664, produced tetrads with two viable and two inviable spores, indicating the presence of a recessive-lethal mutation. We performed array-CGH analysis on JAY664 and determined that two copies of Chr12, including the *ACE2* locus, were present (Fig. S3C-D). Together, these results showed that all five rough colony isolates were homozygous for *ace2-A7*, in agreement with the initial hypothesis that the high frequency of smooth to rough colony morphology transitions among JAY270 derivatives was caused by interhomolog recombination leading to copy-neutral LOH. Unexpectedly, the JAY664 array-CGH also showed that this rough colony isolate did carry copy number alterations in genomic regions other than Chr12. In particular, a terminal deletion on the right arm of Chr6 spanned multiple essential genes and explained the 2:2 spore viability phenotype (Fig. S3E). Interestingly, the breakpoint for the Chr6 deletion occurred at a position immediately distal to *FAB1*, where a tRNA gene and Ty1 retrotransposon sequences are found in the S288c reference genome and in the JAY270 maternal Chr6 homolog, which sustained the deletion. We analyzed the status of HetSNP markers flanking the breakpoint and found that a proximal marker (Chr6 - 185 Kb) remained heterozygous, while a distal marker (Chr6 - 229 Kb) lost heterozygosity through a deletion mechanism (Fig. S4A-B). Even though we did not characterize the precise sequences that were joined at the deletion breakpoint in JAY664, this pattern was consistent with non-allelic homologous recombination (HR) involving Ty retrotransposon repeats, a major class of gross chromosomal rearrangements observed in *S. cerevisiae* (19, 20).

LOH is typically a regional, rather than local, mutational mechanism. Interstitial tracts of homozygosity can span tens of kilobases, and terminal tracts are even longer, extending all the way to the telomeres (21). Therefore, in addition to being homozygous for *ace2-A7*, the rough colony isolates might also be homozygous for flanking HetSNPs. We tested this model initially at low resolution by determining the genotypes at eleven Chr12 HetSNPs using PCR (Table S3). The results of this analysis were compiled to produce the LOH tract maps shown in Fig. 2. As expected, JAY270 was heterozygous for all eleven markers tested. Notably, Chr12 in this strain is only heterozygous for positions to the left of the rDNA cluster (Fig. S1). This pattern is similar to that described previously for other heterozygous diploid *S. cerevisiae* genomes and is suggestive of ancestral LOH events mediated by rDNA instability (7).

**Figure 2.**
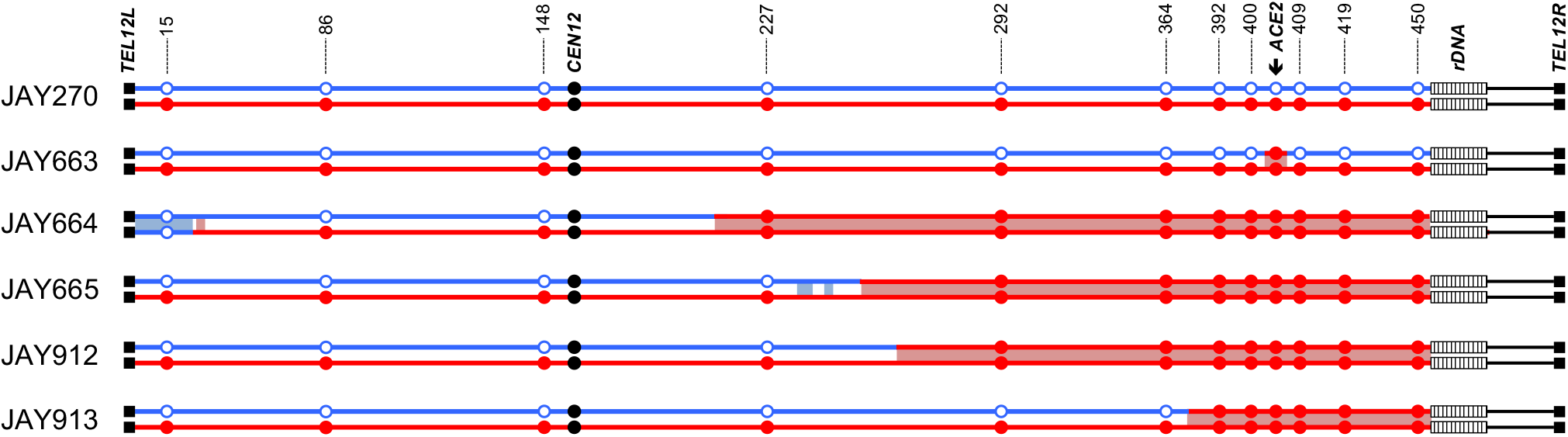
LOH tract maps of Chr12 from five original rough colony isolates. The genotypes at twelve phased JAY270 Chr12 HetSNP marker loci were determined using PCR and RFLP or Sanger sequencing analyses (Table S3). The approximate coordinates of the markers are shown in Kb. The Chr12 homolog containing the *ace2-A7* allele was arbitrarily designated as maternal (Chr12-M, red) and the homolog containing the wild type *ACE2* allele as paternal (Chr12-P, blue). JAY270 was heterozygous at all markers, and all rough colony isolates were homozygous for the *ace2-A7* allele. White boxes distal to the 450 Kb HetSNP represent ~1.5 Mb of ribosomal DNA repeats (rDNA). Chr12 regions distal to the rDNA do not contain any heterozygous markers in JAY270. The red or blue shading corresponds to the directions (M or P, respectively) and approximate breakpoint positions of the LOH tracts determined at high resolution using whole genome sequencing (detailed in Fig. S6 and File S1).

Analysis of the JAY663 isolate showed that, while it was homozygous for the *ace2-A7* mutation, it remained heterozygous at all other flanking markers, including those immediately proximal and immediately distal to the *ACE2* locus. A mitotic gene conversion tract limited to the 8.7 Kb region between these HetSNPs could explain this result. Alternatively, a *de novo* -1 contraction mutation in the adenine homopolymer run of the *ACE2* allele could also account the for the JAY663 genotype (22).

The four remaining isolates (JAY664, JAY665, JAY912, and JAY913) were homozygous for regions well beyond the *ACE2* locus. In this low-resolution map, all four LOH tracts were unidirectional, continuous, and homozygous for SNPs present in the Chr12 homolog that contained the *ace2-A7* allele, which we arbitrarily designated the maternal homolog (Chr12-M; red in all figures). Subsequent high resolution LOH mapping using whole genome sequencing (WGS; below) confirmed the initial results and revealed additional complexities to the tracts. The centromere-proximal breakpoints of the LOH tracts were roughly mapped to positions ranging from 39 Kb (JAY913) to 184 Kb (JAY664) from the *ACE2* locus. On the distal side, these isolates had an additional 45 Kb of LOH that extended to the HetSNP at position 450 Kb, located 1.4 Kb proximal to the rDNA repeats. From this point, Chr12 contains ~1.5 Mb of rDNA repeats plus another ~0.6 Mb of distal homozygous single copy sequences. Since the 450 Kb HetSNP was the most distal marker in Chr12, we could not distinguish if these LOH tracts were generated as very long interstitial gene conversion events, or if they extended to the right telomere. This initial PCR-based analysis also revealed an unexpected secondary LOH event on the left arm of Chr12 in JAY664, but in this case, it was associated with homozygosity for the SNPs from the paternal homolog (Chr12-P, blue in all figures; see below).

### Analysis of selected Chr12 LOH

Taken together, the results described above showed that the majority of the isolates with altered colony morphology were homozygous not only at the *ACE2* locus, but also for surrounding regions, indicating that interhomolog mitotic recombination was frequent in JAY270 and that it likely had substantial effect on the genetic makeup of this strain. In addition, the distribution of HetSNPs in the genome is notably uneven (Fig. S1), with long tracts of homozygosity, suggesting that abundant LOH occurred in the JAY270 lineage.

To gain a deeper understanding of the impact of genome instability processes on the present genetic composition of JAY270, we conducted experiments to directly measure the rate of LOH in this strain (Fig. 3A). Starting with a homozygous *ura3/ura3* derivative of JAY270 (FGY050; gift from F. Galzerani), we introduced one copy of the *KlURA3-ScURA3-KanMX4* CORE2 counter selectable cassette (23) at a position immediately proximal to *ACE2* (1.3 Kb from the adenine homopolymer run). We grew cultures of strains carrying this insertion and plated the cells in media containing 5-FOA to identify clones that had lost the cassette. We performed this assay in a derivative of JAY270 carrying the hemizygous CORE2 insertion in Chr12-M. The frequency of homozygosity for the *ACE2* allele was 1.2 x 10^-4^, comparable to the unselected frequency of *ace2-A7* homozygosity (5 in ~50,000) estimated earlier in the study. We also used hemizygous CORE2 insertions to measure LOH rates at two other positions in the genome (Chr4 near *SSF2*, and Chr13 near *ADH6*), and on Chr5 by deleting one allele of the *CAN1* gene (*can1D::NatMX4/CAN1*), and selecting for loss of the remaining WT allele in clones resistant to canavanine. In order to provide a reference for comparison of LOH rates from JAY270, we introduced these same four constructs in a standard laboratory yeast strain background routinely used to study genome instability mechanisms, including LOH (CG379; Fig. 3B) (24, 25). The rates of LOH were not significantly different between the two strain backgrounds at Chr12 (p=0.225), and were slightly higher in JAY270 at Chr5 (p<0.001) and slightly higher in CG379 at Chr13 (p=0.021) and Chr4 (p<0.001). In general, this analysis indicated that the two strains backgrounds have comparable levels of chromosome stability, and therefore JAY270 genome is not inherently unstable.

**Figure 3.**
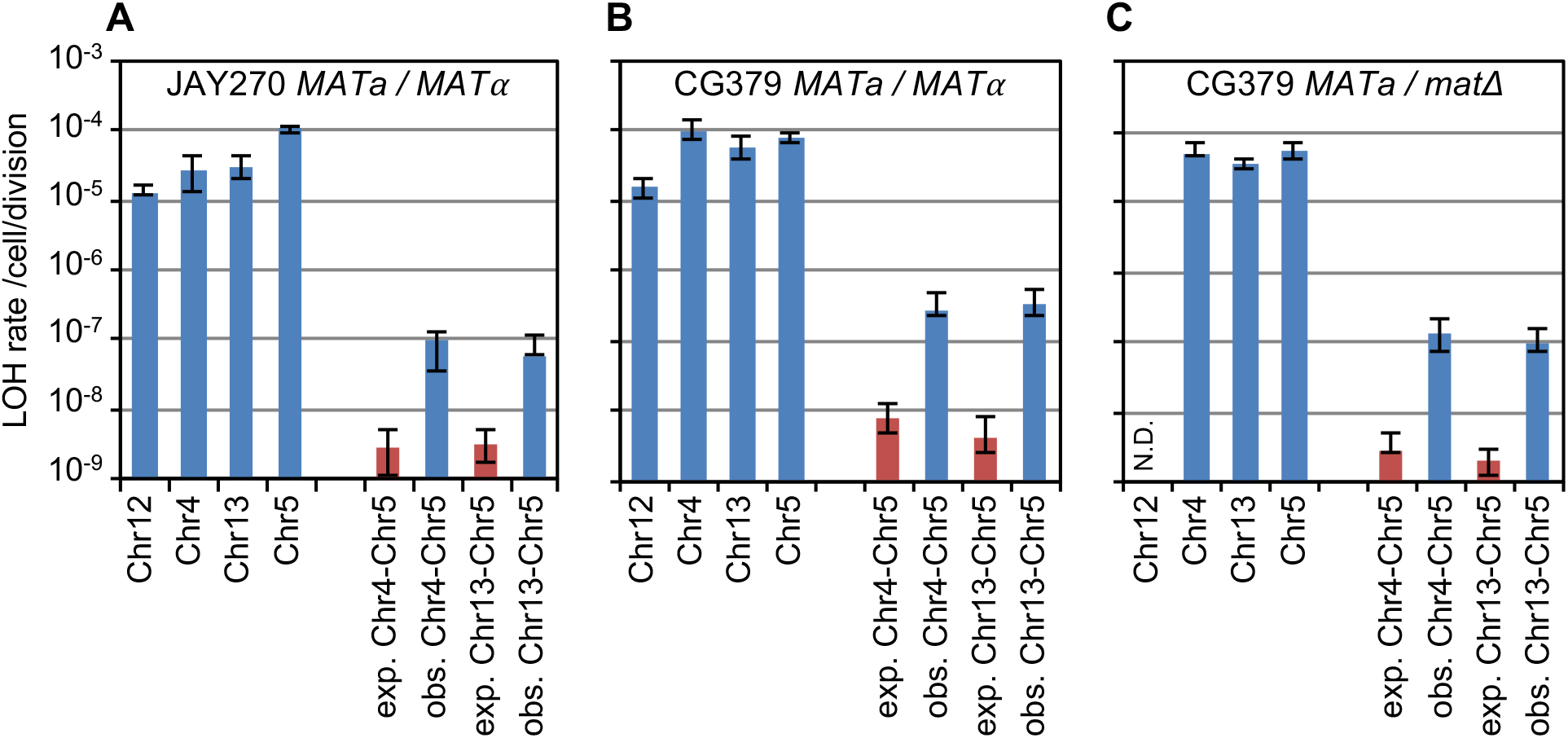
Quantitative analyses of LOH. The measured median rates (blue bars) of single and double LOH events are shown in panels A for the JAY270 strain background, B for the CG379 strain background, and C for CG379 *MATa / matΔ* strains. In the X axis, Chr12, Chr4, or Chr13 indicate diploids with hemizygous insertions of the CORE2 cassette (*KlURA3-ScURA3-KanMX4*) at each of those chromosomes. Chr5 indicates hemizygous deletion of *CAN1*. These strains were used to determine the rates of single LOH individually at each CORE2 insertion or at *CAN1*. Error bars indicate 95% confidence intervals (CI) for each rate measurement. The rates expected for independent double LOH events (red bars; exp.) were calculated by multiplying the respective rates of single LOH at Chr4 and Chr5 or Chr13 and Chr5. The matching rates of observed (obs.) double LOH for each loci pair are shown in blue to the right, measured from strains carrying one copy of CORE2 and one copy of *CAN1*. The expected 95% CI for independent double LOH rates were calculated by multiplying upper or lower 95% CI boundaries from the corresponding single rates, therefore they likely overestimate the expected 95% CI. The rate of single Chr12 LOH for the strain background in panel C was not determined (N.D.).

We also used the 5-FOA^R^ selection approach to characterize the qualitative nature of the Chr12 LOH tracts in a larger set of independent clones (Fig. S5A-B). We used nine PCR-RFLP markers to map LOH tracts from 5-FOA^R^ clones derived from CORE2 insertions on Chr12-M and Chr12-P. The patterns observed were similar between the two LOH directions, and resembled the tracts observed in the five initial spontaneous rough colony isolates (Fig. 2). All 41 selected Chr12 LOH clones were heterozygous for the left arm, and 39 had uninterrupted unidirectional LOH tracts on the right arm, starting at positions between *CEN12* and *ACE2*, and extending up to the 450 Kb HetSNP. This predominant tract pattern was consistent with a simple interhomolog mitotic crossover mechanism.

The region between *CEN12* and *ACE2* was divided in five intervals delimited by HetSNP markers. The distribution of breakpoints found at these intervals was not significantly different between the strains carrying the CORE2 insertion at Chr12-M or Chr12-P (c^2^ = 0.855; *p* = 0.93), suggesting that both homologs shared similar mitotic recombination properties. We pooled the observed breakpoint distribution data from the 19 5-FOA^R^ clones derived from the Chr12-P insertion, and the breakpoints from 23 spontaneous rough isolates that were mapped using WGS (below; Fig. S5C and Fig. S6). We compared the total number of breakpoints leading to *ace2-A7/ace2-A7* LOH observed within each interval to the expected distribution if breakpoints were allocated purely as a function of the size of the physical interval. This analysis indicated that the observed breakpoint distribution was not significantly different from this simple model (c^2^=6.846; *p*=0.1442) (Fig. S5D). Since LOH breakpoints in this region of the genome were relatively evenly distributed, it does not appear that a mitotic recombination hotspot (*i.e.* fragile site) was present.

### Genome-wide analysis LOH and CNA in the JAY664 isolate

In addition to the two copy-neutral Chr12 LOH events and the Chr6 terminal deletion discussed above, we also detected additional structural alterations in the genome of the rough colony isolate JAY664. Using pulse-field gel electrophoresis (PFGE) we detected a size reduction in the long homolog of Chr11 (Fig. 4A), and array-CGH showed a 15 Kb full deletion (0 copies) near the right end of that chromosome. (Fig. 4B). Analysis of PCR markers proximal to (Chr11 - 639 Kb) and within (Chr11 - 653 Kb) the deletion showed that the deleted region was hemizygous in the JAY270 genome (Fig. S4C-E). In JAY664, however, the proximal marker became homozygous, and the distal hemizygous region was lost. The combination of array-CGH and PCR genotyping showed that JAY664 experienced a copy-neutral LOH event with a breakpoint proximal to the Chr11 - 639 Kb marker, leading to homozygosis for the haplotype lacking the hemizygous region represented in the microarray. A similar pattern of proximal LOH and distal CNA was also detected for the right end of Chr7 (Fig. 4C and Fig. S4F-I). Altogether in JAY664, including PFGE, array-CGH and WGS methods, we detected structural alterations at eight independent regions of the genome.

**Figure 4.**
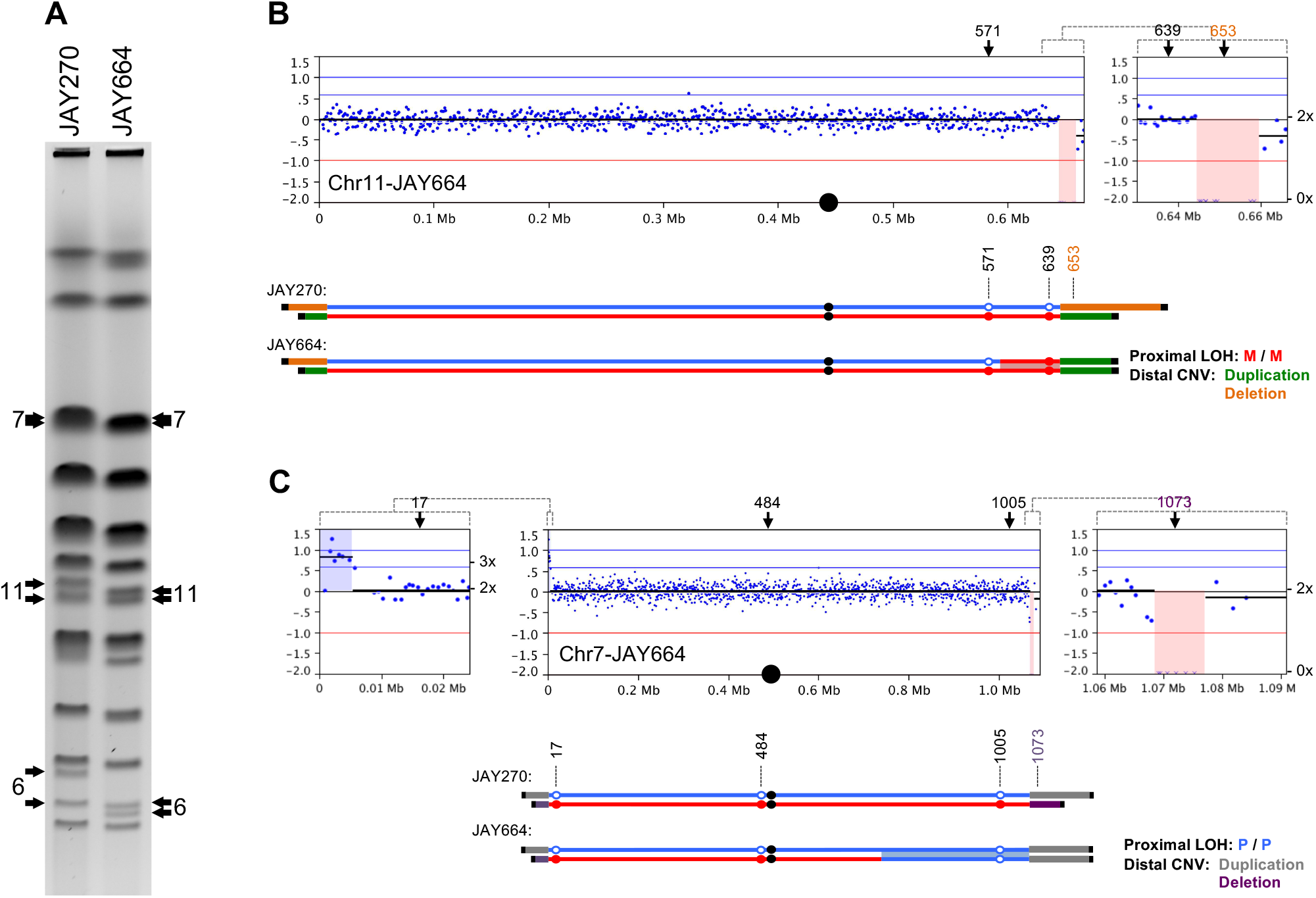
Analysis of unselected chromosomal changes in JAY664 A shows a PFGE of JAY270 and the JAY664 rough colony isolate. Arrows indicate the chromosomal bands that differed between the two strains, with the respective chromosome number next to them. B shows the Chr11 array-CGH copy number plot for JAY664 relative to JAY270. The X-axis indicates the chromosomal coordinates and Y-axis has the Log2 (Cy5-JAY664) / (Cy3-JAY270) ratio, with the corresponding DNA copy number (*i.e.* 0x, 2x, 3x) indicated to the right. Blue dots correspond to individual probes in the array and their Log2 Cy5/Cy3 ratios and positions along the chromosome. The inset to the right shows a close-up view of the right end of Chr11 where a full deletion of the probes in the region (pink shading) was detected. The positions of three PCR markers evaluated in the analysis are indicated, and their genotypes are indicated by blue and red circles according to the presence of paternal or maternal alleles (see Table S3 and Fig. S4). Note that only the sequences from the *S. cerevisiae* S288c reference genome are represented in the array. The probes within the array-CGH deletion signal are hemizygous in JAY270 (orange, 1x), including the 653 Kb PCR marker. JAY664 is homozygous for the maternal right terminal segment of Chr11, and therefore completely lost the signal for the paternal probes in the array, while duplicating maternal JAY270 hemizygous sequences (green). The maternal hemizygous sequences are not present in the S288c genome and thus not represented in the array. The position of the Chr11 LOH breakpoint determined by WGS was near the 581 Kb HetSNP. Panel C shows a similar analysis of Chr7 in JAY664. Gray and purple lines represent the hemizygous regions at the ends of the paternal and maternal homologs, respectively. A proximal LOH event (WGS breakpoint at 784 Kb) and distal CNV event with deletion of one hemizygous region (purple, including the 1073 marker) and duplication of sequences (gray) not present in the array. The inset to the left shows the amplification (blue shading) of probes near the left telomere, but no proximal LOH occurred on that region of Chr7. Instead, the sequences represented by these probes are actually present as a hemizygous insertion near the left telomere of Chr12 in JAY270. An LOH event at that region (see JAY664 Fig. 2 and Fig. S6) caused them to be duplicated, while deleting hemizygous regions not represented in the array.

### Investigation of systemic genomic instability

The remarkably high number of coincident rearrangements observed in JAY664’s genome suggested that they might not have accumulated sequentially or independently. Instead, a simpler scenario would be that the observed karyotype arose during a burst of systemic genomic instability in one or a few ancestor cells in the JAY664 lineage. If this model were correct, then the existence of a primary LOH event should increase the likelihood of existence of secondary LOH events, thus other rough colony isolates with a primary selected Chr12 LOH event should also carry a higher than expected number of unselected LOH events elsewhere in their genomes.

To test this hypothesis, we initially analyzed 29 independent smooth clones derived from JAY270, all isolated after five transfer cycles in liquid culture without single cell bottlenecks (~57 cell generations; Methods). Since these clones were smooth, they should not carry LOH on Chr12, and indeed, PCR analysis confirmed that they were all heterozygous *ACE2/ace2-A7* (data not shown). We performed PFGE to detect unselected chromosomal rearrangements in these isolates lacking a primary LOH event. No visible rearrangements were observed in any of the 29 smooth clones (Figure S7) for chromosomes other than Chr12 (rDNA) and Chr8 (*CUP1* tandem repeats). This observation was corroborated by a second experiment in which we generated two independent mutation accumulation lineages of JAY270 that were cultured with 10 single colony bottlenecks for a total of ~220 cell generations. PFGE analysis of smooth clones from intermediate and final points along the two lineages also showed no visible chromosome size polymorphisms (Fig S8).

The rate of mitotic crossover for the entire yeast genome has been estimated to be ~6.2 x 10^-4^ / cell / cell division (21). Based on this value, we calculated that roughly one of the 29 smooth clones from the liquid growth regimen, and less than one of the two bottleneck lineages, should have had at least one LOH event. Although the PFGE approach can only detect a subset of structural chromosomal changes (interstitial events, LOH in the longer chromosomes and LOH in chromosomes without structural variation between homologs are not detectable), the lack of any visible chromosomal rearrangements in the smooth clones derived from these two experiments indicated that rearrangements were not abundant in JAY270 when a primary LOH event was not selected.

Next we performed a similar liquid growth without bottlenecking regimen analysis, this time plating ~1,000 cells after every passage cycle to identify rough colonies. Twenty independent spontaneous rough colonies were obtained relatively quickly using this approach. Eleven of them were isolated after ~43 or less cell generations, and only one of them was isolated after more than ~57 (JAY1127; ~85 generations, Table S4). PCR analysis of the new rough clones showed that all were homozygous *ace2-A7/ace2-A7* (data not shown). Combined with the original set of five (Fig. 2), a total of 25 independent rough colony isolates were analyzed by PFGE (Fig. 4 and Fig. S9). Strikingly, six of them showed at least one visible size polymorphism for chromosomes other than Chr12 and Chr8. This result showed that, despite its limitations, the PFGE approach was able to detect unselected chromosomal rearrangements in the rough clones, whereas none were seen among the smooth clones. This difference was especially notable since the rough clones were derived from fewer cell generations.

We then used WGS in order to quantify and characterize all LOH events present in the 25 rough colony isolates (Fig. 5). This analysis yielded detailed primary LOH tracts for Chr12 in all clones, and revealed a total of 27 unselected secondary LOH tracts. Based on the estimated rate of genome-wide mitotic crossover (21), we calculated that only one clone out of the 25 should have had one unselected LOH event, in addition to the selected event spanning the *ACE2* locus (6.2 x 10^-4^ crossovers/genome x 57 cell divisions x 25 clones). Note that this was an inflated estimate given that most rough clones analyzed were selected after 43 or less cell divisions (Table S4), and importantly, LOH is only detectable at the ~60% of the JAY270 genome that is heterozygous (Fig. S1). In addition to the high number of total secondary LOH events detected, the distribution of secondary LOH events per clone was also biased. Out of the thirteen clones with unselected LOH, four had two unselected events, three had three unselected events, and finally, one outlier (JAY664) had an astounding seven unselected LOH events. The probability of randomly identifying just one rough clone containing only two unselected LOH events is smaller than 10^-5^. The probability of having the distribution we actually found is far lower. Therefore, our WGS analysis indicated the occurrence of systemic bursts of genomic instability leading to LOH.

**Figure 5.**
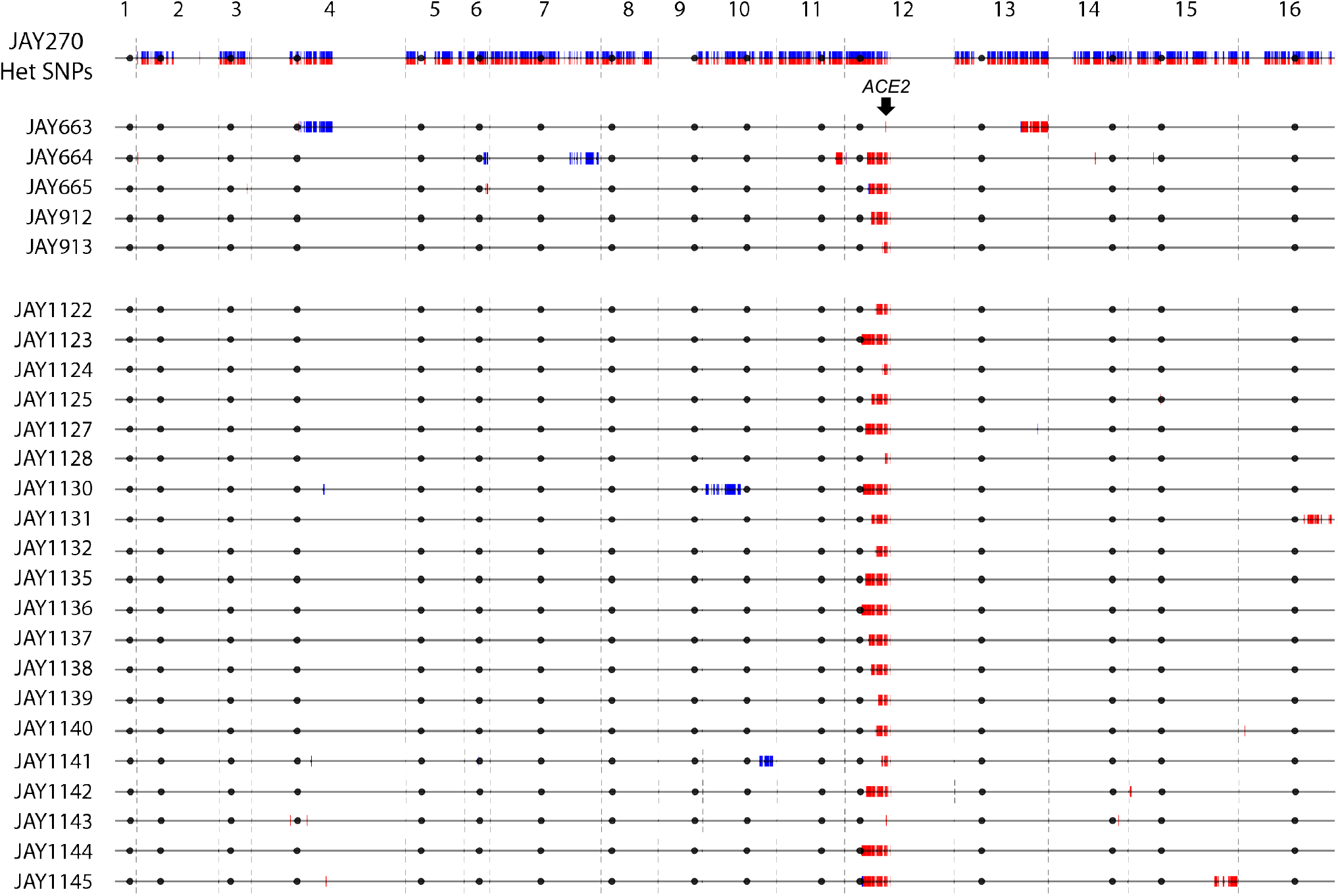
Genome-wide map of LOH tracts in spontaneous rough colony isolates. The top horizontal line is the linear end-to-end depiction of the 16 *S. cerevisiae* chromosomes in JAY270 with HetSNPs represented as paternal (blue) and maternal (red) markers. Chromosome numbers are indicated above, and the position of the *ACE2* locus is shown. Each horizontal line below corresponds to the genomes of the 25 spontaneous rough colony isolates sequenced. Only the HetSNPs that were homozygous P/P or M/M are shown (heterozygous markers are omitted to emphasize visualization of the LOH tracts). As expected from selection for the rough colony morphology, all clones were homozygous for the maternal *ace2-A7* allele (red). In addition, 27 unselected M/M or P/P LOH tracts were detected elsewhere in the genomes of the rough colony isolates. Plots were generated to scale in Python 2.7 using the matplotlib package and a custom script. For size reference, Chr1 is 230 Kb.

The detailed maps of Chr12 LOH in the 25 sequenced rough clones (Fig. S6) showed that 20 tracts were unidirectional, continuous and extended from a position between *CEN12* and the rDNA cluster. In five cases, the tracts had complex discontinuities and some even showed limited LOH for the paternal allele near the breakpoint. Among the unselected LOH tracts (Fig. 5), 15 were interstitial (median size 5.0 Kb) and 12 were terminal (median size 232.1 Kb) consistent with gene conversion and mitotic crossover mechanisms, respectively. We also identified 15 *de novo* point mutations among the sequenced clones. However, the small overall number detected was not sufficient to draw conclusions regarding nucleotide mutational signatures. The full WGS LOH and point mutation analyses in the rough clones is presented in Table S5 and File S1.

### Validation and quantification of coincident LOH

The discovery of multiple LOH tracts in spontaneous rough colonies suggested the possibility that allelic mitotic recombination, and chromosomal rearrangements in general, may arise and accumulate during systemic genomic instability episodes, rather than independently of each other. Alternatively, it could also be possible that homozygosity at or near the Chr12 *ACE2* locus itself could destabilize the genome, thus increasing the likelihood of secondary rearrangements. We conducted a new round of experiments to validate and quantify the observation of systemic genomic instability, and did so under conditions that removed any possible influence from Chr12 status.

Earlier in the study we used diploid strains individually hemizygous for either the counter selectable CORE2 cassette or *CAN1* gene, and measured the rate of LOH at the individual loci by selection for resistance to 5-FOA or canavanine independently. We used the same approach to measure rates of coincident double LOH in diploids that were hemizygous for both CORE2 and *CAN1*, and selected for simultaneous resistance to 5-FOA plus canavanine. If the occurrence of LOH at a CORE2 insertion locus were completely independent from the occurrence of LOH at the *CAN1* locus, then the rate of LOH at both loci should be equal to the multiplicative product of the two individual rates. This independent mutation model predicts very low double LOH rates, in the 10^-9^ /cell/division range (Fig.3A-C, red bars). The double LOH rates that we measured experimentally were 30-100 fold higher than expected. This was observed in two different strain backgrounds (JAY270 and CG379) for double LOH events at two different pairs of genomic regions (Chr4 and Chr5; Chr13 and Chr5), in all cases with Chr12 remaining unaltered.

Finally, we considered the possibility that the systemic genomic instability mechanism causing the high rates of double LOH could result not from a mitotic process, but instead could be due to sporadic and transient initiation of meiotic recombination in a few cells in the population followed by return-to-growth (RTG) (26). Unlike mitotic recombination, which is sporadic and unscheduled, meiotic recombination is initiated as a well coordinated systemic and genome wide event (27). To distinguish between these mechanisms, we repeated the double LOH measurements in diploids deleted for the *MATa* locus. These *MATa/matD* diploids behaved essentially like haploids. They efficiently mated to a *MATa* haploid tester strain and lost the ability to sporulate (data not shown). Since *MATa/matD* diploids are unable to activate the meiotic developmental program, they are also unable initiate meiotic recombination. The rate of single LOH in the *MATa/matD* diploids was similar or slightly lower than the rate in the *MATa/MATa* diploids, consistently with earlier studies (28). Importantly, the observed rate of double LOH was ~50 fold higher than expected if the single LOH events were initiated independently of each other. Taken together, our results support the existence of a mitotic systemic genomic instability (mitSGI) process in yeast cells, which is independent and qualitatively different from sporadic initiation of meiosis and RTG.

## DISCUSSION

### Rough colony phenotype and mutation in *ACE2*

We showed that a colony morphology transition in JAY270 resulted from an LOH event at a region of Chr12 heterozygous for a mutation in the *ACE2* gene. This gene encodes a transcription factor that controls the expression of genes involved in septum destruction and mother-daughter cell separation (17). The *ace2-A7* -1 frameshift allele found in JAY270 leads to a premature stop codon resulting in a truncated, likely inactive, protein that lacks three zinc finger domains and a nuclear localization sequence (29). Interestingly, another heterozygous diploid industrial strain, FostersB used in brewing (8), has a +1 frameshift variant in the same adenine homopolymer region of *ACE2* (*ace2-A9* allele). The *ACE2* gene has also been shown to be involved in the transition between the yeast and hyphal forms of *C. albicans*, an important trait for pathogenesis. In that context, inactivation of the *ACE2* ortholog contributes to the formation of cell filaments during hyphal growth, and an alternative isoform of the Ace2 protein helps prevent inappropriate activation of cell detachment from hyphae (30). In *S. cerevisiae*, and in particular in JAY270 during bioethanol production, it is not known if or how the switch between dispersed to aggregated cell growth states has an effect on fitness. However, it is possible that JAY270’s heterozygous *ACE2/ace-A7* genotype may offer an advantage by giving the population the ability to quickly access the aggregated state through LOH. This may provide a short-term adaptive solution as cells encounter various environmental challenges during industrial fermentation (9). Individuals in the population could then return to the dispersed state through expansion or contraction of the adenine run (22), or through sporulation and mating to an *ACE2* haploid.

### mitSGI and precedents of coincident recombination

Beyond the genetic characterization of the rough colony phenotype in JAY270, this study allowed us to uncover the mitSGI phenomenon through which multiple LOH events can accumulate in a cell lineage. Using PFGE, array-CGH and WGS, we determined that yeast clones carrying a primary selected LOH tract at Chr12 were more likely than expected to carry unselected LOH tracts. We also showed in quantitative LOH assays that combinations of double LOH at Chr5 and Chr4 or Chr13, occurred at rates 30-100 fold higher than expected if single LOH events occurred independently. We interpret these results as evidence for the occurrence of bursts of genomic instability leading to multiple LOH events over one or few mitotic cell generations.

Spontaneous mitotic recombination events like the ones described here are triggered by local DNA lesions and/or replication fork collapse episodes, which then lead to chromosomal breakage and allelic HR repair using the homolog as template (31). Such precursor lesions are thought to occur randomly in vegetative cells, both spatially and temporally, therefore mitotic recombination events involving different chromosomes should not be coincident. In contrast, meiotic recombination is known to be a systemic genetic variation process, since it occurs simultaneously throughout the genome and involves intricate coordination between generation and repair of genome wide double strand breaks (27).

While our study is, to our knowledge, the first to describe the mitSGI phenomenon through the lens of high-resolution genome-wide analytical methods, there have been sparse reports of elevated coincident mitotic recombination in yeasts as well as in mammalian cells dating back decades (32–39). The typical experimental design in those cases was to select clones for carrying a recombination event at a primary locus, and then screening the resulting clones for the occurrence of secondary unselected recombination at one or a limited number of unlinked loci. The same intriguing observation, shared in all cases, was a frequency of coincident recombination that was higher than that predicted assuming the individual events occurred independently.

In some of the yeast studies, the high coincident recombination rates were interpreted as being derived from a small number of cells within the replicating population that spuriously entered the meiotic developmental program, or transiently experienced a “para-meiotic” state, but reverted back to mitotic growth (33, 34). A recent study specifically characterized this type of return-to-growth (RTG) event and the genome-wide recombination outcomes associated with it (26). The authors often detected a large number of LOH tracts per clone (minimum of 5, average of ~30, and up to 87), indicating that the RTG induction leads to abundant and widespread recombination. Another notable finding was that while interstitial LOH (gene conversion-like; GC) tracts were frequent, their sizes were relatively constrained (2.3 Kb on average). This measurement is notable because it is consistent with GC tract sizes measured in haploids derived from complete meiotic divisions; ~2 Kb median size (40). In contrast, GC tracts associated with mitotic recombination tend to be significantly longer, approximately 5-6 Kb median size (21). This variation in typical GC tract sizes is likely a reflection of subtle mechanistic differences in the processing of HR intermediates between meiotic and mitotic cells.

Despite the possibility discussed above, there are several reports of high coincident recombination in proliferating cells in which the induction of a full meiotic cycle, RTG or para-meiosis were either unlikely or could be ruled out entirely. One study in *S. cerevisiae* specifically measured the formation of spurious haploids from mitotic diploid cultures displaying high coincident intragenic recombination at unlinked pairs of heteroalleles (36). The authors found that while haploids did form in their cultures, the frequency was far below that needed to influence the formation of double recombinants, thus concluding that a low level of cryptic meiosis was not a likely contributor. In addition, one of the seminal studies (37) of LOH in *C. albicans* (a species devoid of a conventional sexual cycle (5), reported data that closely parallel our own observations. First, the authors selected clones for the presence of a primary LOH event at the *GAL1* locus on chromosome 1. Then, using a low resolution SNP-array platform, they detected frequent unselected secondary LOH tracts among clones carrying the primary event, but rarely in control clones still heterozygous at *GAL1*. In addition, selection for LOH at *GAL1* was associated with the emergence of altered colony morphology phenotypes, presumably derived from rearrangements elsewhere in the genome. Accordingly, clones displaying altered morphology were enriched for the presence of unselected LOH tracts when compared to clones with normal morphology.

Another important pair of precedents of mitSGI observations comes from experiments conducted in mammalian systems. These used either human TK6 lymphoblastoid cells in culture (38), or mouse kidney cells *in vivo* and in culture (39). In both cases, the starting cells were heterozygous for mutations at the counter-selectable markers, *TK* and *Aprt*, respectively, enabling the selection of clones carrying a primary LOH event at those loci. Subsequently, the presence of secondary LOH tracts was assessed at roughly a dozen loci elsewhere in the human or mouse genomes. The two studies found that secondary LOH was more frequent in clones selected for carrying the primary LOH event than in controls clones that remained heterozygous. These studies demonstrated that mitSGI also exists in metazoans, and can be detected in cells that are exclusively mitotic, thus ruling out a contribution from meiotic recombination, at least in these contexts.

The studies outlined above suggest that cryptic initiation of meiosis in a small number of cells can in some cases lead to systemic genomic instability, however, we favor the interpretation that the events analyzed in our study originated primarily from *bona fide* mitotic cells. The recent work by Laureau *et al*. (26) clearly defined the features of systemic LOH caused by meiotic initiation followed by RTG. The pattern we detected in our study was different, and instead was consistent with mitotic patterns. The number of unselected interstitial LOH (GC) tracts per clone we detected was small (typically 1 or 2) and their sizes were long (median 5.0 Kb). This was reinforced by the observation that *MAT*a/*mat*Ddiploids, incapable of entering the meiotic developmental program, continued to display double LOH rates that were far higher than expected from independent events.

### mitSGI-like observations in human disease

In addition to the experimental examples above, our results also resemble recent reports of bursts of mitotic genomic instability in humans during cancer genome evolution and early development. Specifically, genome-wide copy number profiling of thousands of individual cells isolated from tumors in 12 patients with triple-negative breast cancer revealed that a large number of CNAs were acquired within a short period of time at the early stages of tumor development (15). Most of these CNAs were shared between several cells from a same tumor, suggesting the occurrence of a burst of genomic instability in one or few initiating cells followed by a long period of stable clonal expansion. Although the study had power to detect gradual accumulation of mutations, no clones with intermediate CNA profiles were identified, suggesting a punctuated model of mutation accumulation.

Another pertinent parallel is the recent analysis of patients with genomic disorders that carry multiple *de novo* constitutional CNVs (MdnCNVs; (16). Typically in those patients, only one of the structural variants was the primary event causing the symptoms associated with the disorder. The additional CNVs were secondary, occurred at unrelated regions, and apparently formed during a short burst of genomic instability at some point in the perizygotic time interval. The changes then propagated stably during development to be found in all cells in the patients. Taken together, these results suggest that mitSGI processes may be universal and may play an important role in human disease development.

### Possible mechanisms underlying mitSGI

Our results so far suggest that mitSGI is not likely associated with initiation of meiotic recombination and RTG. However, the specific causes for the existence of a small subset of recombination-prone cells within a normal mitotic population remain to be determined. This phenomenon could well have multiple and distinct origins, however, we favor two non-exclusive mechanisms, related to cellular ageing and stochastic gene expression. These two models are attractive because they are transient in nature, which would support stable transmission of rearranged genomic structures after the systemic vulnerability time window has passed.

The first scenario is that clones carrying multiple unselected LOH events originated from replicatively old mother cells. This model stems from the observation of a marked increase in the rate of LOH in daughter yeast cells budded from mothers that had undergone ~25 cell divisions (41), relatively old within the context of a maximum *S. cerevisiae* replicative lifespan of ~40. Subsequent work from the same group showed that this increase in nuclear genomic instability was strongly correlated with the initial appearance of mitochondrial DNA loss and/or damage in the old mother cells (42). In our study, however, all of the spontaneous rough colony isolates analyzed retained normal respiratory activity (all were non-*petite*; grew on non-fermentable carbon sources), so they must have had integral mitochondrial genomes. They also did not show signs of continual genomic instability. Therefore, if replicative aging were an underlying factor in mitSGI, it would be through a pathway that does not involve loss of mitochondrial function.

Another explanation for a subpopulation of hyper-recombinogenic cells involves heterogeneities that exist even within an isogenic population. Specifically, cell-to-cell variation (*i.e.* noise) in gene expression has been reported in organisms ranging from prokaryotes, to yeast, to humans (43). It is plausible that stochastic variation in the expression of a broad class of genes involved in genome stability could cause specific protein levels to drop below those required for optimal function. A recent comprehensive genome stability network analysis identified 182 genes involved in suppression of gross chromosomal rearrangements (44), and an earlier genetic screen identified 61 genes specifically involved in suppressing LOH (45). In this scenario, rare individual cells that fail to adequately express any of these genes could effectively behave as null mutants for a short period. Some of these genes act cooperatively, therefore concomitant loss of activity causes extreme levels of genomic instability. For example, double knockouts for *TEL1* and *MEC1*, encoding critically important DNA damage response proteins (orthologs of mammalian ATM and ATR, respectively), show marked increase in mitotic genomic instability (46), often accumulating multiple genome rearrangements (47). A similar extreme phenotype might be expected in a wild type cell that by chance simultaneously had a critically low level of transcription of two genome stability genes. Likewise, overexpression of single genes encoding a subunit of a genome stability multi-protein complex could lead to a dominant negative phenotype that temporarily impairs function. Importantly, the hyper-recombinogenic state of these individuals would be completely reversible once the descendant cells returned to the gene expression levels typical of most individuals in the population. This mechanism could explain the observed stable clonal expansion that followed mitSGI in our spontaneous rough colony clones, as well as in the recent *in vivo* human studies (15, 16).

The analysis of spontaneous rough morphology clones provided unprecedented detailed information about the nature and frequency of secondary recombination events resulting from the mitSGI process. Our study also provides a unifying context for the interpretation of classic and recent reports of coincident recombination in yeasts, in mammalian experimental systems, and in human disease. The combination of whole genome analyses and the double LOH selection approach described here offer a powerful experimental platform to further dissect the core mechanisms responsible for the mitSGI phenomenon.

## MATERIALS AND METHODS

An extended description of all experimental procedures used is available in *SI Materials and Methods*. A summary of the methods is provided below.

### Yeast genetic backgrounds, growth media and procedures

*Saccharomyces cerevisiae* strains used in this study descended from either the JAY270 or CG379 strain backgrounds (Table S1). Standard procedures for yeast transformation, crossing and sporulation were followed (48). Cells were grown in YPD and synthetic minimal media (SC). *URA3, natMX* and *hphMX* transformants were selected in uracil drop-out SC, YPD plus 200 mg/L of nourseothricin (Nat) or 300 mg/L of hygromicin, respectively. Counter-selection against *URA3* and *CAN1* were performed in SC plus 1g/L of 5-FOA and 60 ml/L of canavanine in arginine drop-out, respectively.

### Isolation of spontaneous rough colonies derived from JAY270

The five spontaneous rough colony isolates described early in the study were visually identified in YPD plates containing diluted liquid cultures of JAY270, or JAY270 isogenic diploids. The precise number of passages in liquid media until isolation of these clones is unknown. The 20 independent rough colonies obtained later were selected during liquid YPD growth passages, in which independent JAY270 cultures were plated daily for visual screening of colonies. The growth cycle in which spontaneous rough colony isolates for each culture were identified is provided in Table S4. 29 random control smooth isolates were selected after 5 consecutive growth cycles.

### Quantitative LOH rate assays

Single colonies were inoculated into 5 ml liquid YPD, and incubated for 24 hours at 30^o^ C in a rotating drum. The cultures were serially diluted and plated on YPD (permissive), and SC plus 5-FOA (selective) and/or canavanine (selective). Colony counts were used to calculate recombination rates and 95% confidence intervals using the Lea & Coulson method of the median within the FALCOR web application [http://www.keshavsingh.org/protocols/FALCOR.html] (49, 50). Statistical analyses of pairwise comparisons between recombination rates were performed using a two-sided nonparametric Mann Whitney test in GraphPad Prism software.

### Genome Sequencing Analyses

The Illumina short read WGS platform was used to sequence the genomes of the JAY270 parent strain, the 25 rough colony isolates and 56 JAY270-derived haploids. The sequencing reads were used to build a phased draft map of HetSNPs in JAY270’s genome, to map the *ace2-A7* mutation and to detect the LOH tracts in the rough colony isolates. All genome sequencing data associated with this study is available in the Sequence Read Archive (SRA) database under study number SRP082524.

## ACKNOWLEDGMENTS

We thank Tom Petes, Michael McMurray, Dmitry Gordenin, and Aaron Mitchell valuable insights and/or comments on the manuscript. NMVS received a pre-doctoral fellowship from Brazil’s CAPES (0316/13-0). PC and AVP were supported by a UGC fellowship. KTN was supported by a Wellcome Trust-DBT India Alliance Intermediate fellowship- (IA/I/11/2500268) and IISER-TVM intramural funds. Research reported here was supported by the NIH award number R35GM119788 to JLA.

